# Sand fly synthetic sex-aggregation pheromone co-located with insecticide reduces canine Leishmania infantum infection incidence: a stratified cluster randomised trial

**DOI:** 10.1101/712174

**Authors:** Orin Courtenay, Erin Dilger, Leo A. Calvo-Bado, Lidija Kravar-Garde, Vicky Carter, Melissa J. Bell, Graziella B. Alves, Raquel Goncalves, Muhammad M. Makhdoomi, Mikel A. González, Caris M. Nunes, Daniel P. Bray, Reginaldo P. Brazil, James G. C. Hamilton

## Abstract

**Objective:** To evaluate the efficacy of a synthetic sex-aggregation pheromone of the sand fly vector *Lu. longipalpis*, co-located with residual insecticide, to reduce the infection incidence of *Leishmania infantum* in the canine reservoir, and to reduce sand fly vector abundance. To compare the outcomes to those resulting from fitting deltamethrin-impregnated collars to the canine reservoir.

**Methods:** A stratified cluster-randomised trial was designed to detect a 50% reduction in canine incident infection after 24 months in 42 recruited clusters, randomly assigned to one of three intervention arms (14 cluster each): pheromone + insecticide, insecticide-impregnated dog collars, or placebo control. Infection incidence was measured by seroconversion to anti-*Leishmania* antibody, *Leishmania* parasite detection and canine tissue parasite loads. Changes in relative *Lu. longipalpis* abundance within households were measured by setting three CDC light traps per household.

**Results:** A total 1,454 seronegative dogs were follow-up for a median 15.2 (95% C.I.s: 14.6, 16.2) months per cluster. The pheromone + insecticide intervention provided 13% (95% C.I. 0%, 44.0%) protection against anti-*Leishmania* antibody seroconversion, 52% (95% C.I. 6.2%, 74·9%) against parasite infection, reduced tissue parasite loads by 53% (95% C.I. 5.4%, 76.7%), and reduced household female sand fly abundance by 49% (95% C.I. 8.2%, 71.3%). Variation in the efficacy against seroconversion varied between trial strata. Equivalent protection attributed to the impregnated-collars were 36% (95% C.I. 14.4%, 51.8%), 23% (95% C.I. 0%, 57·5%), 48% (95% C.I. 0%, 73.4%) and 43% (95% C.I. 0%, 67.9%), respectively. Comparison of the outcomes of the two interventions showed no statistically consistent differences in their efficacies; however, the errors were broad for all outcomes. Reductions in sand fly numbers were predominant where insecticide was located (chicken and dog sleeping sites), with no evidence of insecticide-induced repellency onto humans or dogs.

**Conclusion:** The synthetic pheromone lure-and-kill approach provides protection particularly against *L. infantum* parasite transmission and sand fly vector abundance. The effect estimates are not dissimilar to those of the insecticide-impregnated collars, which are documented to reduce canine infection incidence, and human infection and clinical VL disease incidence, in different global regions. As a low-cost alternative or complimentary vector control tool, optimisation of best community deployment of the pheromone + insecticide are now underway.

**Author summary:** The sand fly vector of the intracellular parasite *Leishmania infantum* causing human and canine visceral leishmaniasis in the Americas is *Lutzomyia longipalpis*. Dogs are the proven reservoir. Vector control tools to reduce transmission suited to this predominantly exophilic vector are lacking. Insecticide-impregnated dog collars protect dogs against infectious bites from sand fly vectors, resulting in reductions of new infections in both dogs and humans. However, collars are costly particularly for endemic communities, and alternative approaches are needed. Recent bulk synthesis of a sex-aggregation pheromone produced by male *Lu. longipalpis* was shown to attract large numbers of conspecific females to lethal pyrethroid insecticides. This study, conducted in Brazil, evaluated the efficacy of this novel lure-and-kill approach to reduce seroconversion and infection incidence with *L. infantum* in the canine reservoir, in addition to measuring its impact on household abundance of *Lu. longipalpis*. Deployed in 14 stratified clusters, the outcomes were compared to those resulting attributed to the collars fitted to dogs in another 14 clusters; each intervention was compared to the 14 clusters that received placebo treatments. The beneficial effects of the lure-and-kill method were most noticeable on confirmed infection incidence and clinical parasite loads, and in reducing sand fly abundance. The overall effect of the two interventions were not statistically dissimilar, though the confidence intervals were broad. We conclude that the novel low-cost lure-and-kill approach should be added to the vector control toolbox against visceral leishmaniasis in the Americas.

## Introduction

Sustainable control of arthropod vectors to reduce infectious disease transmission represents a major challenge confronting public health programmes[1]. Standard approaches such as indoor residual spraying of insecticides (IRS) or insecticide treated nets (ITNs), are most effective against insecticide-susceptible vector populations that are endophilic and/or bite when hosts at risk are under ITNs e.g. as against *Anopheles gambiae*[2]. Suboptimal insecticide-based vector control occurs when contact rates with insecticide treated surfaces by susceptible vectors is less frequent[3], as expected following IRS/ITNs against exophillic vectors. One potential solution is to lure biting vectors to strategically placed insecticide using attractant semiochemicals (kairomones and pheromones). Specific insect pheromones mediate conspecific mating (sex), aggregation, oviposition or invitation behaviour[4]. In the agricultural sector, integrated pest management programs deploy pest pheromones to monitor and reduce pest populations and disrupt pest mating aggregations, with the aim to limit crop yield loss, environmental damage, and insecticide use[4-6]. In contrast, whilst some pheromones produced by vectors of public or veterinary health importance have been identified e.g.[7], they appear to be absent or not characterised in many of the most important human and animal disease vectors. Indeed, to our knowledge, there are no published studies that have tested the efficacy of a vector pheromone to reduce infection or disease incidence.

One important vector species that produces a large amount of sex-aggregation pheromone is *Lutzomyia longipalpis* (Diptera: Psychodidae). This is the principal vector of *Leishmania infantum* (Kinetoplastida: Trypanosomatidae) in the Americas, a protist parasite that causes human and canine zoonotic visceral leishmaniasis (ZVL)[8]. Domestic dogs are the proven reservoir host[9], though non-reservoir (“dead-end”) hosts, such as chickens and other domestic livestock, are significant blood sources that are assumed to maintain sand fly populations[10].

The majority of incident human ZVL cases occur in Brazil[8], where the national ZVL control program include human case detection and treatment, and IRS of houses and animal sheds within 200m of a newly detected case[11, 12]. To reduce the canine reservoir population, the program recommends test-and-slaughter or chemotherapeutic treatment of *Leishmania* infected dogs, canine vaccination and/or application of topical insecticides[11]. Despite this extensive arsenal of control tools, there is no apparent decline in human case incidence[13-15]. On the contrary, ZVL has geographical expanded into new regions and into urban settings[15-17]. Thus, sustainable alternative or complimentary methods to combat transmission are still needed.

The recent bulk synthesis of the male *Lu. longipalpis* sex-aggregation pheromone[18] provides such an opportunity[19]. Male *Lu. longipalpis* release the pheromone from abdominal glands that attracts conspecific males and appetitive females. The resulting leks are formed on or near animal hosts, where the sand flies copulate and the females blood-feed, which results in *L. infantum* transmission[20-23]. In field experiments, the synthetic pheromone attracts significantly more *Lu. longipalpis* to experimental chicken sheds than to those without the synthetic pheromone[24]. And when co-located with pyrethroid insecticide applied to experimental sheds, it attracts and kills significantly more *Lu. longipalpis* compared to untreated control sheds[25]. In a long-lasting controlled release formulation, the pheromone is attractive for up to 3 months[19]. Field trials are now needed to evaluate the efficacy of this novel lure-and-kill approach to reduce *Leishmania* transmission.

Here we report the results of a stratified cluster randomised trial, conducted in Brazil, to test the efficacy of the synthetic pheromone co-located with a pyrethroid insecticide, to reduce (i) the incidence of *Leishmania* exposure and infection in the canine reservoir; (ii) the abundance of *Lu. longipalpis* around households; and (iii) to compare these outcomes in relation to parallel deployment of deltamethrin-impregnated Scalibor collars fitted to dogs.

## Methods

### Study location

The study was conducted between July 2012 and May 2016 in semi-urban towns and in suburban districts of Araçatuba city (21204011S; 50458883W), located in the administrative region of Araçatuba, N.W. São Paulo state, Brazil (Figure 1). ZVL expanded within the region over the last two decades, where it is now considered endemic[16, 26-29]. The human case incidence was 6.3 per 100,000 with a case fatality rate of 9% recorded in 2011 just prior to the study[27]. This represents the highest human VL incidence within São Paulo state which recorded 2,332 autochthonous cases between 1999 and 2013[28, 30]. Canine seroprevalences in the study region ranged from 12-45% (Superintendência de Controle de Endemias [SUCEN], unpublished data).

**Figure 1.** Map of study region in Aracatuba, NW São Paulo state, administrative district. Spatial distribution of trial clusters under control (○) collar (△) and pheromone (⬒) interventions. Magnified section for clusters in Araçatuba city.

### Study design

The trial was designed as a stratified cluster randomised trial (CRT) where the towns (municipalities), and Araçatuba subdistricts, were designated as independent clusters. Clusters, households and dogs were selected in a three– step procedure (Figure 2).

**Figure 2.** Study design and structure.

### Recruitment

#### Clusters

Forty towns within the Araçatuba administrative region and 12 subdistricts of Araçatuba city, were listed for potential trial inclusion as independent intervention clusters. Cluster inclusion criteria included (i) evidence of recent transmission (at least one confirmed human or/and canine infection) within the 4 years prior to the intervention study by inspection of human case records[30], and canine testing records in 2006-2008 (SUCEN, unpublished data). (ii) That the location was within feasible driving time (1.5 hours) of the trial operations centre in Araçatuba; and (iii) that each cluster was geographically distinct, separated by ≥1 km to minimise any inter-cluster contamination by dispersing *Lu. longipalpis* sand fly vectors. Mark-release-recapture studies show ≥97% of *Lu. longipalpis* recaptures are within 300m of the release location[21, 31, 32].

Thirty-three municipalities and 9 districts of Araçatuba (42 clusters in total) met these inclusion criteria (Figure 2), being located within an area of c. 11,250km^2^ (Figure 1).

#### Houses and dogs

Local health authorities provided lists of households within clusters for potential recruitment based on inclusion criteria that (i) the household maintained at least one chicken (dead-end host) and at least one seronegative dog (defined below) at the time of recruitment; and (ii) the householder(s) and their animals were normally resident. Following consultation with local health authorities, and written permission provided by the municipality health officer, informed written consent was obtained from dog owners to test their dog(s) for anti-*Leishmania* antibodies (described below).

### Cluster stratification

Clusters were then stratified, each assigned to one of three strata based on the initial pre-intervention canine seroprevalence within the cluster, of >50% (strata 1: “high” n=18 clusters), <50% (strata 2: “low” n=15 clusters), or Araçatuba location (strata 3: “mixed high and low” n=9 clusters) (Figure 2). Araçatuba clusters were placed in a separate strata as being the regional capital it was considered to have better resources to manage ZVL, knowledge, attitudes and practises (KAP) characteristics, and demographics that could affect transmission dynamics in ways different to in the towns[33].

### Randomisation and treatment allocation

Clusters received one of three treatments, namely (i) synthetic pheromone lure co-located with pyrethroid insecticide; (ii) pyrethroid-impregnated collar fitted to dogs, or (iii) placebo control. These are described below. Within the three defined strata, clusters were ranked in descending order of pre-intervention seroprevalence, and then randomly assigned to one of the three interventions by random number generator in STATA software. All subsequent within-stratum triplet clusters were similarly assigned alternately to intervention groups, resulting in 14 clusters in each treatment arm.

### The Interventions

#### Synthetic pheromone lures and insecticide arm

The synthetic pheromone formulation (*±*-9-methylgermacrene-B [CAS RN: 183158-38-5]) was a copy of the (S)-9-methylgermacrene-B pheromone produced male *Lu. longipalpis*. 10mg of the pheromone was sealed in an 8 cm × 3 cm polythene sachet prototype dispenser designed for slow release (Russell-IPM Ltd. UK), and equivalent to natural pheromone release by 80,000 male *Lu. longipalpis* over a 3 month period[19]. Each household received a lure placed within 1m of the main chicken roosting site. Co-located with the pheromone, micro-encapsulated lambda-cyhalothrin (LC-ME) (®Demand 2.5cs, Syngenta, Brazil) was applied at 20mg a.i. m^-2^ to surfaces close to chicken roosting sites using a GUARANY 441-10 compression sprayer (Guarany Industriae Comercio Ltda, Itu, SP). Sprayed sites included (i) all available surfaces in and on chicken coops (32.6% of sites), (ii) from ground level up to 3m of the roosting tree paying special attention to roosting branches (52.5%), or (iii) 3m^2^ (1.5m x 2m) wall surfaces next to ground perches (7.7%), or similar unusual sites (7.2%). Pheromone lures and insecticide were replaced on 9 occasions at an average interval of 91 (S.D. 20.0) days (Supplementary S1).

#### Insecticide-impregnated collar arm

Scalibor collars (Intervet, MSD Saúde Animal, São Paulo, Brazil) impregnated with 40mg g^-1^ deltamethrin, consisted of a 65cm white polyvinyl chloride strip weighing 25g. Collars were fitted to dogs ≥3 months old following the manufacturer’s instructions. All seronegative dogs per houses were fitted with a Scalibor collar at baseline. At subsequent recruitment rounds, newly eligible dogs were fitted with a collar irrespective of serological status. The veterinary team fitted and replaced any lost collars, promoted their correct use to dog owners, and recorded any adverse reactions and reasons for collar losses. Scalibor collars have an activity period of 5-6 months against sand flies according to the product label, though longer durations of >10 months are experimentally demonstrated[34-36]. Collars were routinely replaced on 5 occasions at an average interval of 182 (S.D. 12.1) days (Supplementary S1).

#### Control arm

All dogs in control clusters received a placebo collar at recruitment, houses received an identical lure dispenser containing no pheromone, and chicken roosting sites were sprayed with water. For logistic reasons, spraying was limited to at baseline and 368 days later (Supplementary S1), whereas newly eligible dogs were fitted with a placebo collar. Losses of placebo collars or lures were replaced at subsequent dog recruitment rounds (Supplementary S2).

### Blood sampling and sampling regime

Peripheral blood was collected from dogs by veterinarians by venepuncture onto two replicate Whatman 3MM Chromatography papers (GE Healthcare, 3030-614), which were then dried at ambient temperature, labelled, placed in individual zip plastic bags containing silica gel (Geejay Chemicals), and stored at 4°C until processing. Sera eluted from the filter papers were tested for anti-*Leishmania* antibodies by ELISA. Up to 5mls of blood was also collected into EDTA tubes to harvest leukocytes for molecular detection of *L. infantum* parasites by quantitative PCR (qPCR). The laboratory procedures are described in Supplementary S3.

At baseline clinical examination, veterinarians scored dogs for eight non-specific signs of canine leishmaniasis: alopecia, dermatitis, hyperkeratitis, skin lesions, conjunctivitis, onychogryphosis (excessive nail growth), lunettes, uveitis, and lymphadenopathy (enlarged popliteal lymph nodes). Each sign was scored on a semi-quantitative scale from 0 (absent) to 3 (severe), or 0 to 2 (for onychogryphosis and hyperkeratitis). Scores were then summed to give an overall clinical severity score.

Canine blood samples were collected at baseline to identify potential recruits, and recruited dogs then sampled again at their final follow-up.

### Trial outcome measures

The primary outcome measure was cluster-level cumulative seroconversion incidence in naïve dogs in each intervention arm compared to in the control arm. The secondary canine outcome measures were *L. infantum* parasitological infection cumulative incidence, and changes in blood parasite loads (*L. infantum* genome equivalents per ml blood buffy coat), confirmed by specific qPCR. The third outcome measure was relative changes in cluster-level household counts of *Lu. longipalpis* measured by setting CDC miniature light traps as described below.

Canine clinical condition is not considered a reliable marker of infection incidence following others[37]; signs of canine leishmaniasis are non-specific, thus positive diagnosis requires extensive differential diagnosis, and statistical power was insufficient in this study to rely on changes in advanced canine VL disease.

### Sand fly catches

After the final canine follow-up sample in October 2015, the trial interventions as described above were continued following the same regime for an additional 7 months until the end of May 2016. Sand fly sampling was conducted in 40 of the 42 clusters in 6 approximate quarterly trapping rounds from January 2015 to May 2016 (January/February; April; July/August; October/November in 2015; and January/February; and April/May in 2016). Data from 2 clusters were incomplete and thus excluded from the analyses. The final dataset was generated from 209, 188 and 193 (n=590) trapping nights in 129, 121 and 113 (n=363) houses, in 14, 13, and 13 control, pheromone + insecticide and collar intervention clusters, respectively.

Miniature CDC light traps with the light bulb removed were positioned in three locations per household: inside the house, at the dog sleeping site or kennel entrance, and at the principal chicken roosting site, each trap associated with the respective host: humans, dogs and chickens. Captured sand flies were sexed and counted under a stereomicroscope. Specific identification was not performed as the vast majority (>98%) of peridomestic sand flies captured in the study locations during parallel entomological studies were confirmed to be *Lu. longipalpis* by dissection of spermathecae[38].

### Data collection

Veterinary staff collected details of the dog’s history, and information on any newly acquired dogs, host numbers, and losses of dogs, collars and pheromone lures, by verbal questionnaire to household heads by house-to-house visitation and/or by active phone contact at least every 3 months.

### Sample size calculations

The sample size was calculated for the primary outcome measure (canine seroconversion incidence) for a two-treatment randomised control trial[39, 40], whereby cluster-level outcomes in collar and in pheromone treated clusters were each compared to the outcomes in the control clusters.

The trial was statistically powered to detect a 50% reduction in *L. infantum* seroconversion incidence in naïve dogs after 24 months follow-up with a baseline canine instantaneous annual incidence of 0.6, estimated from canine testing between 2006-2008 in the region (SUCEN, unpublished data) (see Table 2). Calculations were based on a surveyed harmonic mean of 24 dogs per cluster (assuming 1 negative dog recruit per household), with equal numbers of clusters per arm, and coefficient of variation between clusters κ_m_=0.40. The latter value was more conservative than κ_m_=0.34 estimated from the variation in cluster-level canine seroprevalences (1,701 dogs in 42 clusters) at initial trial recruitment. Under these design conditions, 12 clusters per arm were required per intervention arm to achieve a statistical power of 90% with 95% confidence to reject the null hypothesis.

**Table 1.** Summary of the total dogs sampled, recruited and with follow-up sample, according to the trial intervention arm to which the dogs were subsequently allocated.

| Intervention arm | Dogs initially sampled for potential recruitment | Seronegative dogs recruited (proportion of sampled) | Negative dogs with follow-up sample (proportion of recruits) |
| --- | --- | --- | --- |
| Control | 1,575 | 961 (0.610) | 455 (0.473) |
| Pheromone | 1,702 | 994 (0.584) | 480 (0.483) |
| Collar | 1,641 | 1,016 (0.619) | 519 (0.411) |
| Total | 4,918 | 2,971 (0.604) | 1,454 (0.489) |

**Table 2.** Serological infection estimates for dogs surveyed prior to trial recruitment, and canine historical testing records in the same region.

| Sample period | Population sample | FOI estimates<sup>1</sup> |  | Seroprevalence<sup>2</sup><br>(pos/n dogs) |
| --- | --- | --- | --- | --- |
| | | $\lambda$ incidence/year<br>(SD), n dogs | $\rho$ recovery/year<br>(SD) | |
| 2012-2014<sup>a</sup> | Control arm | 1.14 (0.163), 1,558 | 1.48 (0.264) | 0.610<br>(961/1575) |
|  | Pheromone arm | 1.47 (0.225), 1,693 | 1.82 (0.335) | 0.584<br>(994/1702) |
|  | Collar arm | 1.14 (0.159), 1,631 | 1.52 (0.265) | 0.619<br>(1016/1641) |
|  | Trial arms combined | 1.26 (0.151), 4,882 | 1.64 (0.235) | 0.604<br>(2,971/4,918) |
| 2006-2008<sup>b</sup> | 19 regional towns | 0.61 (0.094), 2,304 | 0.63 (0.121) | 0.422<br>(972/2,304) |
| 2006-2008<sup>c</sup> | 20 regional towns |  |  | 0.451<br>(4592/10,186) |

<sup>1</sup> FOI estimates were calculated from fitting age-seroprevalence data to an incidence ( $\lambda$ )-recovery ( $\rho$ ) model.
<sup>2</sup> Crude seroprevalences estimated including dogs without age records.
<sup>a</sup> estimates calculated from dogs pre-recruitment resident in the 14 towns (= trial clusters) per intervention arm to which they were subsequently randomly allocated; serological infection detected using the ELISA described in this study.
<sup>b</sup> FOI estimates calculated from canine samples accompanied by dog age data; serological infection detected using an ELISA kit (EIE-CVL, Bio Manguinhos/Fiocruz-RG, Brazil);
<sup>c</sup> estimates for the same regional population as in (<sup>b</sup>) but including also test data without dog age records.

To buffer effects of potential loss-to-follow-up (LTF) of clusters and dogs, the number of clusters enrolled was increased to 14 clusters per arm. By the end of the trial no LTF of clusters occurred, though substantial LTF of dogs and houses occurred (see Results). Recalculations at the end of the study, revealed a harmonic mean 35 dogs followed-up for an average 16.4 months per cluster, with a starting instantaneous incidence of 1.26 year^-1^ measured from age-prevalence data in all recruited dogs prior to being under intervention (Table 2). From these data, the trial design provided 90% statistical power to detect an equivalent 44% reduction in infection incidence between trial arms.

### Statistical analysis

Analysis of the intervention effect on seroconversion and parasite detection incidence were computed using mixed effects binomial complimentary log–log models expressed as incident risk ratios (IRR). Random intercepts for trial clusters were fitted (trial cluster being the higher level of structuring in the data[41], and log_10_ normalised days under intervention set as the model offset. Similarly, negative binomial mixed effects models were used to test the intervention effects on log_10_ +1 transformed *Leishmania* parasite loads (ml^-1^), and on sand fly numbers.

Complimentary log–log model fits were achieved by Gauss–Hermite numerical adaptive quadrature of the random-effects estimators (quadchk routine in STATA), validated using 16 integration points in model run comparisons to confirm quadrature fitting accuracy; model runs showed ≤0.01% variation in resulting estimates, thus considered to be reliable[42].

For infection outcomes, models comprised variables included *a priori* on the basis that they could affect the trial balance, namely strata (3-levels), baseline canine exposure (baseline anti-*Leishmania* antibody titre), proportion of time (days) under intervention in the bimodally high (December-May) (*cf*. low June - November) sand fly season[43], and household mean numbers of dogs and chickens as surrogates of host odour intensity that may invoke a density-dependent or competing attractant[20, 31, 44]. For analyses of sand fly data, *a priori* covariates included strata, sand fly trapping period (6 levels) reflecting sand fly seasonality, and host abundance at the time of trapping i.e. numbers of people, dogs and chickens associated with each of the 3 trapping locations per household. The model incorporated a cluster term for trial clusters.

The outcomes from these models were considered “unadjusted” effect estimates. Unadjusted estimates were then adjusted on detection of significant model improvement by individual inclusion of additional demographic variables in the model, namely, predominant chicken roosting site category (described above); month and sand fly season (as above) of dog recruitment; dog age at recruitment [median: 24m; IQR: 8-48m)]; dog sex; property type [house or small holding]; general clinical condition score of dog at recruitment [median: 3.1; IQR: 2-4)]. Each variable was evaluated for significance by log–likelihood ratio test (LRT) of nested models.

In a secondary analysis, the outcome × strata (3-level) interaction term was tested against the full model to evaluate differential intervention effects between trial strata. In the case that the interaction term was significant with ≥90% probability, individual strata-level effect estimates were further investigated.

The equality of variances in cluster-level dog follow-up time (days) was tested using the Levene’s robust test statistic adapted by Brown & Forsythe[45] to provide robust estimators of central tendency (median [W50] and 10% trimmed mean [W10]).

Data were analysed in STATA v.15 (StataCorp LP, College Station, TX).

### Force of Infection model estimation

The instantaneous incidence (force of infection, FOI) was calculated for dogs serologically tested prior to trial recruitment, by fitting the age-prevalence data to a standard age-incidence-recovery model[46, 47].

To evaluate changes in infection rates from c. 6-8 years earlier, the FOI was similarly calculated using historical data with age records for 2,304 dogs resident in 19 towns in the same region, sampled in 2006-2008 (SUCEN, unpublished data). Positives were identified on detection of “*L. major*-like” promastigote soluble antigens[48] by an ELISA-based kit (EIE-leishmaniose-visceral-canina-Bio-Manguinhos [EIE-LVC], Bio-Manguinhos/Fiocruz-RJ, Brazil). Seroprevalence in the same historical populations was estimated by inclusion of an additional 7,882 dog results that did not have age records (n=10,186 dogs in total) (see Table 2).

### Data management and masking

Diagnostic results and household questionnaire data were entered into data-checking entry forms designed in ACCESS 2007 relational database by a trained technician, and databases then checked for inconsistencies. Unblinding for final analysis was conducted independently after all dogs had been tested by laboratory staff who were blinded to the cluster treatments and to cluster and household identities through a bar-coding system; all tested sample tubes were bar-coded and results subsequently matched to dog ID bar-codes in the database.

### Ethical considerations

The trial protocols for dogs were approved by the Committee for Ethical Use of Animals (CEUA [FOA-00124-2013]), UNESP, Brazil, and the Animal Welfare and Ethical Approval Body (AWERB, [48723]), University of Warwick, UK. Household questionnaire designs were approval by the Biomedical and Scientific Research Ethics Committee (BSREC, REGO-2015-1388), University of Warwick, UK. Informed written consent was obtained from dog owners to sample and fit collars to their dogs, and from the town and district health authorities to conduct the study within their administrative jurisdiction.

## RESULTS

### Pre-enrolment canine infection estimates

A total 4,918 dogs were serologically tested prior to trial recruitment, of which 2,971 dogs (60%) were seropositive (Table 2). Seroprevalence and FOI estimates were similar between dogs in the three trial arms to which they were subsequently allocated (Table 2; Figure 3). No statistical differences were detected in these infection measures between the trial arms, accounting for the trial structure, date of recruitment, dog age and trial cluster (mecloglog mixed effects model: z<0.89, *P*>0.38). Notably these pre-intervention infection rates were higher than equivalent estimates calculated from canine serosurvey records in the same region conducted a number of years previous (Table 2; Figure 3).

**Figure 3.** Canine age-seroprevalence data (symbols) fitted to an incidence-recovery model to provide the best fit (lines) from which annual FOI (incidence l and recovery r) were estimated (results shown in Table 2). Data include 4,882 resident dogs in 42 trial clusters sampled prior to recruitment, categorised here according to the intervention arm to which the clusters were subsequently randomised: pheromone (⬒, **----**, n=1693), collar (◊, **……**, n=1631), and control (○, **- - -** n=1558) arm. Data also shown for 2,304 dogs resident in 19 towns in the same region sampled in 2006-2008 (△, solid line).

### Dog recruitment and follow-up

The initial enrolment included 630 seronegative dogs under intervention by July-November 2012 (Supplementary S2). The study experienced substantial LTF of dogs and houses primarily due to dogs being lost through mortality or unknown causes and/or households no longer maintaining chickens or eligible dogs. Thus, to fulfil the statistical power requirements, new dogs and houses were recruited between November 2012 and October 2014 (Supplementary S2). This resulted in a total 2,971 seronegative dogs recruited and placed under the trial interventions, of which, 1,454 (48.9%) dogs, resident in 789 houses across the 42 trial clusters, remained in the study for follow-up testing (Table 1). A median 1 (95% C.I.: 1, 2) seronegative dogs was enrolled per house which did not differ between treatment arms (Poisson: z<0.34, *P*>0.16).

The follow-up cluster population was observed for a *per capita* median 17.1 (95% C.I.s: 15.2, 17.7, n=455 dogs), 14.7 (95% C.I.s: 14.0, 16.7, n=480), and 15.2 (95% C.I.s: 13.2, 15.5, n=519) months under control, pheromone and collar interventions, respectively (Supplementary S4). Similar fractions of the follow-up times (0.507, 0.454 and 0.416) fell within the seasonally high period of sand fly abundance (December to May) (LRT: χ^2^_(2)_ =1.08, *P*=0·58). The variance in cluster-level dog follow-up days were not dissimilar between intervention arms (Levene’s W_10_ [df: 2,39]=1.78, *P*=0.838; W_50_ [df: 2,39]=1.60, *P*=0.853). The epidemiological data for the three intervention arms (Tables 1, Figure 3), indicated that the randomization process achieved good trial balance.

### Intervention outcomes

#### Seroconversion incidence

Of the seronegative recruits, 225 (49.5%), 217 (45.2%), and 182 (35.1%) in control, pheromone and collar arms respectively, seroconverted by the end of the study. The annual cluster seroconversion incidence varied between the three trial strata within intervention arms (Figure 4; Supplementary S4).

**Figure 4.** Mean annual incidence (+/-SD) of seroconversion (grey bars) and confirmed parasitological infection (black bars), between trial strata (1 to 3) within control, pheromone and collar intervention arms.

Accounting for the variables describing the trial structure and follow-up intervals, the unadjusted seroconversion incident risk ratio (IRR) was 0·88 (95% C.I. 0.66, 1.16) in the pheromone arm, and IRR= 0.65 (95% C.I. 0.48, 0.87) in the collar arm, each compared to the control arm (model fit: Wald χ^2^ (8)=31·4, *P*<0.0001) (Table 2). The protection attributed to the interventions were therefore 12% (95% C.I. 0%, 33.8%), and 35% (95% C.I. 13.2%, 51.8%) respectively.

Potential adjustment to these estimates was assessed by inclusion of additional demographic variables in the model. Only one significant covariate was identified: the location of the chicken principal roosting site i.e. where the pheromone + insecticide was co-located (LRT: χ^2^ (3) =9·57, *P*=0·023). This led to slight modifications of the effect estimates (Table 2).

**Table 2.** Seroconversion incident rate ratio (IRR) amongst dogs under each intervention: unadjusted and adjusted estimates calculated relative to the control arm, by fitting to mixed-effect complimentary log-log models.

| | Unadjusted IRR<br>(95% C.I.s), n dogs | $P \leq$ | Adjusted<sup>1</sup> IRR<br>(95% C.I.s), n dogs | $P \leq$ |
| --- | --- | --- | --- | --- |
| **(A) Strata 1-3** | N=1,443 |  | N=1,322 |  |
| Pheromone | 0.88 (0.662, 1.159) | 0.353 | 0.87 (0.660, 1.152) | 0.320 |
| Collar | 0.65 (0.482, 0.868) | 0.004 | 0.64 (0.482, 0.856) | 0.003 |

<sup>1</sup> effect modification adjusted by variables significant in LR test: chicken predominant roosting site where pheromone and insecticide were co-located.

Insecticide treatment of the two most common roosting site categories, low trees (53%), and chicken shelters (33%), were not dissimilar in intervention effect; the significant modification was associated with the third roosting site category, most commonly hollows in the ground, but which represented only 7% of all roost sites. In the latter case insecticide was sprayed on the nearest wall.

Neither intervention resulted in significant changes in the mean log_10_ anti-*Leishmania* antibody units in seroconverted dogs (z<1.45, *P*>0.10).

In a secondary analysis, potential differences in the intervention effects on seroconversion incidence between trial strata were examined. This provided evidence of strata-level variation (strata × treatment interactions, LRT test: χ^2^ _(4)_ =8.34, *P*=0.078), observed only in the pheromone arm, suggesting a negative impact in Aracatuba city (stratum 3, n= 3 clusters) (IRR=1.66 [95% C.I.s: 0.971, 2.849], *P*=0.064) (pheromone arm × stratum 3 interaction: z=2.36, *P*=0.018). In contrast, the pheromone effects in strata 1 and 2 (n=11 town clusters) suggested a protective effect of 27% (95% C.I.s: 0.02%, 46.8%) (IRR=0.73 [0.532, 0.998], *P*=0.048).

### Parasitology

Buffy coat samples from 775 recruited dogs at follow-up were tested for the presence of *Leishmania* kDNA in peripheral blood by qPCR. Parasites were detected in 117 (15.1%) of dogs overall; including 20.8% (64/308) of dogs that seroconverted and 11.4% (53/467) of dogs that did not. The latter category of dogs was neither differentially associated with the date or season of recruitment, or their follow-up time, to suggest a predominance of prepatent dogs in the recruited sample.

The percent reduction in the crude number of parasite positive dogs attributed to pheromone and collar interventions at follow-up were 43.3% and 26.1%, respectively (Table 3). Accounting for the trial structure, follow-up periods and covariates as described above, the respective reductions in confirmed *Leishmania* infection incidence were 51.5% (95% C.I. 6.2%, 74·9%) and 22.5% (95% C.I. 0%, 57·5%) (Table 3). The intervention outcomes did not significantly vary between strata (test of treatment × strata interaction term: LRT: χ^2^_(4)_ =4.10, *P*=0·393), nor were effect modifications significant by inclusion of additional demographic variables (LRT: χ2(3) <2.68, *P*>0·444).

**Table 3.** Confirmed *Leishmania* infection incidence, tissue parasite loads, and intervention effects in recruited dogs at follow-up.

| Intervention arm | qPCR<sup>1</sup> positive/dogs tested (%) | parasite infection incidence IRR (95% C.I.s) | geometric mean<sup>2</sup> (95% C.I.) parasite load ml<sup>-1</sup> | parasite load ml<sup>-1</sup> IRR (95% C.I.s) |
| --- | --- | --- | --- | --- |
| control | 48/231 (20.8) | referent | 18.6 (7.18-47.96) | referent |
| pheromone | 25/246 (10.2) | 0.485 (0.251, 0.938) $P=0.032$ | 6.48 (0.85-49.35) | 0.469 (0.233, 0.946) $P=0.034$ |
| collars | 44/298(14.8) | 0.775 (0.425,<br>1.141) $P=0.404$ | 4.82 (1.72 -<br>13.54) | 0.524 (0.266,<br>1.03) $P=0.062$ |
| Total | 117/775<br>(15.1) |  |  |  |

<sup>1</sup> quantification of *Leishmania* kDNA in canine peripheral blood leukocytes per ml<sup>-1</sup> by qPCR, standardised to the endogenous control.
<sup>2</sup> GM calculated in 117 positive dogs with qPCR counts

#### Leishmania parasite loads

The geometric mean *Leishmania* parasite loads per ml^-1^ in the 117 qPCR positive dogs were highly variable (Table 3). Relative to control clusters, the intervention reduced parasite loads in the pheromone arm by an average 53.1% (95% C.I. 5.4%, 76.7%), and in the collar arm by an average 47.6% (95% C.I. 0%, 73.4%) (Table 3). The intervention effects did not significantly vary between strata (test of treatment × strata interaction term: LRT: χ2(4) =0.02, p=0.905), nor were reductions in parasite loads related to the number of days under intervention (LRT: χ^2^_(4)_ =0.39, *P*=0.532), or modified by inclusion of additional demographic variables (LRT: χ^2^_(3)_ <0.21, *P*>0·967).

For the 308 dogs that seroconverted with parasite counts, the log10 parasite loads were not correlated with corresponding log10 IgG antibody units (Spearman’s r=0·074, *P*=0.20); similar non-significant patterns were observed across treatment arms. For the dogs that failed to seroconvert, there were no associations between the fraction that were parasite positive and the date of recruitment, or between their log10 follow-up time (fully adjusted model: z>0.049, *P*>0.23). The annual incidence of confirmed parasitological infection and seroconversion post intervention were also not correlated (Figure 5).

**Figure 5.** Association between annual incidence estimates of confirmed parasite infection and seroconversion at follow-up in control (○) collar (△) and pheromone (⬒) intervention clusters.

### Sand fly abundance

Complete sand fly trapping records were available for 590 trap nights in 363 houses in 40 trial clusters (Supplementary S5). The number of trap nights (trapping effort) were similar between intervention arms (t<1.33, *P*>0.19) and between trial strata (t<1.57, *P*>0.124). Relatively few *Lu. longipalpis* were captured per house, and only 46% of houses were positive for sand fly capture; which was similar under each intervention (Supplementary S5).

The pheromone intervention significantly reduced the numbers of female and male sand flies captured at households relative to controls, whereas the collar intervention tended to reduce only the number of females (Table 5). No consistent differences in sand fly captures were observed between the trial strata (test of intervention arm × strata interaction term: z<1.15, *P*>0.248). Inclusion of additional demographic variables did not significantly modify these effect estimates.

**Table 5.** Intervention effects^1^ on household numbers of male and female *Lu. longipalpis* sand flies captured across 3 CDC lights traps per house.

| Treatment arm | Female sand flies |  | Males sand flies |  |
| --- | --- | --- | --- | --- |
| | IRR (95% C.I.s) | $P \leq$ | IRR (95% C.I.s) | $P \leq$ |
| Pheromone | 0.51 (0.287, 0.918) | 0.03 | 0.44 (0.200, 0.974) | 0.04 |
| Collar | 0.57 (0.321, 1.007) | 0.05 | 0.94 (0.460, 1.918) | 0.86 |

<sup>1</sup> estimated from negative binomial mixed effects models including *a priori* predictors.

### Changes in the distributions of vectors at households

Changes in sand fly numbers in CDC traps placed at human, dog and chicken sleeping sites were further examined. In placebo clusters, the majority of *Lu. longipalpis* were captured at chicken sleeping sites, with fewer but similar numbers associated with dog sleeping sites, and humans (i.e. inside houses) (Table 6).

**Table 6:** Distribution of captured *Lu. longipalpis* at households^1^.

| Sand flies | Arm | No. on people<br>(proportion of<br>treatment total) | No. on dogs<br>(proportion of<br>treatment<br>total) | No. on<br>chickens<br>(proportion of<br>treatment<br>total) | No.<br>total |
| --- | --- | --- | --- | --- | --- |
| Females | Control | 34 (0.23) | 43 (0.29) | 72 (0.48) | 149 |
|  | Pheromone | 20 (0.26) | 28 (0.37) | 28 (0.37) | 76 |
|  | Collar | 19 (0.22) | 17 (0.20) | 49 (0.58) | 85 |
| Males | Control | 45 (0.16) | 52 (0.18) | 188 (0.66) | 285 |
|  | Pheromone | 31 (0.25) | 46 (0.37) | 46 (0.37) | 123 |
|  | Collar | 47 (0.15) | 37 (0.12) | 228 (0.73) | 312 |
| All | Control | 79 (0.18) | 95 (0.22) | 260 (0.60) | 434 |
|  | Pheromone | 51 (0.26) | 74 (0.37) | 74 (0.37) | 199 |
|  | Collar | 66 (0.17) | 54 (0.14) | 277 (0.70) | 397 |

<sup>1</sup> In each household, a single CDC light trap (excluding the bulb) was set inside the house (humans), and outside the house above the sleeping dog, and at the main chicken roosting site.

In the pheromone arm, reductions of 66% (95% C.I. 36%, 81.7%) and 69% (95% C.I. 43.6%, 82.6%) were observed in female and male sand flies captured at the chicken roosting site, being the site of pheromone + insecticide co-location (Table 7). In the collar arm, there was a mean 52% (95% C.I. 0%, 87.9%) reduction in female sand flies at dog trapping sites attributed to collars, although this narrowly failed to reach statistical significance (Table 7). These reductions were not mirrored by significant changes in sand fly numbers at the corresponding alternative trap locations (Table 7).

**Table 7.** Intervention effects on *Lu. longipalpis* abundance at host-associated CDC trap sites

| Capture site | humans |  | dogs |  | chickens |  |
| --- | --- | --- | --- | --- | --- | --- |
| | IRR (95% C. I.s) | $P \leq$ | IRR (95% C. I.s) | $P \leq$ | IRR (95% C. I.s) | $P \leq$ |
| **Females** |  |  |  |  |  |  |
| Pheromone | 0.72 (0.285, 1.814) | 0.49 | 0.71 (0.385, 1.304) | 0.27 | 0.34 (0.183, 0.640) | 0.001 |
| Collar | 0.55 (0.268, 1.115) | 0.10 | 0.48 (0.221, 1.052) | 0.07 | 0.58 (0.296, 1.152) | 0.12 |
| **Males** |  |  |  |  |  |  |
| Pheromone | 1.12 (0.451, 2.758) | 0.81 | 0.80 (0.329, 1.950) | 0.62 | 0.31 (0.174, 0.564) | 0.001 |
| Collar | 0.93 (0.446, 1.957) | 0.86 | 0.76 (0.347, 1.641) | 0.48 | 1.15 (0.564, 2.447) | 0.71 |

Males made up the majority of captures which was not unexpected (Table 6). The number of female sand flies was positively associated with the number of male flies in the same trap (z=7.24, *P*<0.0001), but not with the prevailing mean numbers of household chickens or dogs (z<0.49, *P*>0.26). This relationship was not dissimilar across intervention arms (test of intervention arm × male fly number interaction term: z<0.197, *P*>0.53).

### Comparison of the synthetic pheromone versus collar intervention effects

Direct statistical comparisons of the pheromone *versus* collar intervention outcomes (i.e. not compared to the control arm), did not provide evidence of substantial differences between the two interventions. Only in analysis of seroconversion incidence did collars provide an apparent 25.3% (95% C. I.s: 1.2%, 43.4%) additional protection over the pheromone intervention (IRR = 0.747 [0.566, 0.988], *P*=0.041). Whereas, the pheromone intervention resulted in a 49% (11%, 66.4%) greater reduction in male *Lu. longipalpis* at households compared to in the collar arm (IRR = 0.51 [0.336, 0.790], *P*=0.002). Otherwise, no other statistical differences were detected.

## Discussion

The synthetic pheromone intervention reduced the incidence of confirmatory parasitological infection by 52%, and the geometric mean peripheral blood parasite loads by 53%. The same intervention also reduced the household numbers of female *Lu. longipalpis* by 49%. These promising outcomes were not mirrored in changes in seroconversion incidence across all trial strata. In the 11 semi-urban town clusters (strata 1 & 2) under this intervention, seroconversion was reduced by an average 27%, whereas among the three clusters in Araçatuba city (stratum 3), seroconversion incidence was increased rather than decreased. The latter result was specifically attributed to a single Araçatuba treated cluster, in which the annual seroconversion incidence was 0.0274, which was 2.2× the average (0.0126/year) for the three Araçatuba control clusters. The equivalent rates in the other two pheromone-treated clusters (0.0142 and 0.0139/year) were similar to that of controls (Supplementary S4). There was no further evidence of significant variation in intervention effects between strata in either the pheromone or collar arm. By design, all nine clusters recruited within the regional capital Araçatuba, were assigned to a separate stratum based on the perceived enhanced ZVL control activities and conditions in the city[33]. Compliance may have been lower in this stratum, but we did not detect significant differences in dog recruitment, the LTF of dogs, collars, or pheromone lures, or their replacement rates (data not shown).

### Collars

In contrast, the deltamethrin-impregnated collar intervention reduced canine seroconversion incidence by 36% (95% C.I. 14.4%, 51.8%). The attributable reductions in tissue parasite loads of 48% (95% C.I. 0%, 73.4%), and in household female sand flies of 43% (95% C.I. 0%, 67.9%), were indicative, though failed to reach statistical significance (*P*<0.062).

The collective studies of Scalibor collars in Brazil[49-53], Europe[54-57], North Africa[58], and central Asia[59], demonstrate their impact on *L. infantum* transmission, providing a median 56% (IQR: 48.9%-85.9%; range: 46.9%-100%) protection against canine seroinfection incidence. Community-wide collar interventions provided 43%-50% protection against human infantile seroconversion and clinical ZVL incidence[48, 59, 60]. Furthermore, reductions in *Lu. longipalpis* household abundance[61], and *Lu. longipalpis* infection rates with *L. infantum*[49] are associated with collar interventions. Thus, in the current study, the collar intervention arm acted as a positive control for the previously untested synthetic pheromone lure-and-kill trial method.

Despite this, the overall protection provided by the collar arm appeared somewhat inferior to that of the pheromone arm, when each was compared to the control arm. However, a consistent difference in their performance was not detected by direct statistical comparison of their effect estimates.

### Household sand fly distributions

In control households, the majority of sand flies were captured at chicken roosting sites compared to at dog sleeping sites and inside houses. The pheromone intervention diminished female and male sand flies at chicken roosting sites by 49% and 56% respectively. The collar intervention reduced female, but not male, *Lu. longipalpis* numbers at dog sleeping locations by 43%. There was no evidence that sand flies were diverted from the treated trap sites to the two alternative untreated host trap sites within households. These results are consistent with the insecticidal effects of the associated interventions including the purposeful co-location of insecticide at chicken roosting sites.

Possible imprecision in the effect estimates arises if the synthetic pheromone recruited additional numbers of sand flies that circumvented traps e.g. through insecticide-induced knockdown or excito-repellency. In this case, it is likely that effect estimates reported here are an underestimate of the true intervention effect. Further experiments are needed to quantify these mechanisms.

## Implications for ZVL control

ZVL control guidelines in Brazil recommends IRS of houses, but also of animal shelters[11], where the majority of *Lu. longipalpis* are typically captured[21, 22, 62]. Field studies in north Brazil show sand fly numbers in animals shelters to decrease more or less immediately after insecticide application, but, in parallel, with colonisation of nearby unsprayed sites e.g. household dining huts[63]. Evidenced by additional data[20], the authors of those studies proposed that this shift in vector distribution is a partial consequence of insecticide-induced mortality of male flies causing a decline in pheromone release and recruitment to treated sites, making untreated colonised sites more attractive. Based on this rationale therefore, the co-location of synthetic pheromone should maintain sand fly recruitment to insecticide-treated sites. Results supporting this hypothesis demonstrate that the synthetic pheromone can “restore” female and male recruitment to recently sprayed sheds[25]. Indeed, the synthetic pheromone lure tested in this study attracts approximately 24 times more *Lu. longipalpis* to chicken sheds compared to sheds without synthetic pheromone[19]. This novel lure-and-kill approach offers a potential improvement to the standard IRS practise against ZVL.

A single 10mg lure synthetic pheromone lure is active for 10-12 weeks[19], and attracts *Lu. longipalpis* over distances of 30m (Gonzalez et al., unpublished data), sufficient to cover the typical urban and more rural household vicinity. The attraction of female *Lu. longipalpis* is pheromone dose-dependent[23], in line with density-dependent associations between female and males numbers observed in natural leks / CDC light trap catches (this study; [64, 65]). Chickens are a dead-end (sink) host for *Leishmania*[10], and often the most abundant domestic host in endemic regions[33], hence an obvious location to place the pheromone and insecticide. However, the need for synergistic effects between the synthetic pheromone and host odour to attract *Lu. longipalpis*[66] no longer appears critical as confirmed by on-going field studies, and opening the door for its wider deployment at households without animal hosts. Community-wide experiments are now needed to optimise pheromone doses in different demographic settings.

With respect to transmission dynamics, of particular relevance is the observed reductions in tissue *L. infantum* parasite loads. This is expected to diminish the canine population’s infectiousness to *Lu. longipalpis*, since canine skin, blood, and bone marrow *L. infantum* loads correlate with the dogs’ ability to transmit *L. infantum* to *Lu. longipalpis*[37, 67-70]. Dogs with the highest parasite loads are responsible for the majority of transmission events[37, 68], though not exclusively so[71]. Related to this is the proposal that variation in *Leishmania* metacyclic inoculum from sand fly bites contribute to an individual host’s infection pathology, and subsequent onward transmission potential[72, 73]. Additional supporting data (Bell et al., unpublished data) show Scalibor collar protection to effectively lower the over-dispersion in canine population antibody responses to *Lu. longipalpis* salivary antigens delivered by sand fly bites, and indicative of the level of biting exposure. In simple terms, this predicts that a lower fraction of dogs would receive high density *L. infantum* challenge under this intervention.

The suggested increase in canine FOI from 2006-2008 to the trial period (Table 2) is supported by independent reports of the rise and spread of canine infection, and *Lu. longipalpis* abundance, across São Paulo state[13, 16]. Together with current trends in human ZVL burdens in Brazil[13-15], the need for sustainable vector control is clear. The MoH policy of culling seropositive dogs continues to be unpopular amongst dog owners[74], and in the Brazilian setting, dog collars, along with the registered canine vaccine Leish-Tec, and anti-*Leishmania* chemotherapy treatment options, may be too costly and/or perceived insufficiently effective to achieve community-level compliance[75, 76]. Scalibor collar labels indicate 5-6 months effective duration, though collar losses from dogs are variably high (range: 0.6-8.2% per month)[50, 52-55, 57, 59, 60], necessitating replacement, particularly in regions of year-round transmission. In this trial, collar losses were 7.5% (95% C.I.: 6.5, 8.6) per month, compared to pheromone lure loss of 2.7% (95% C.I.: 0.26, 5.2) per month.

## Conclusions

Manipulation of vector behaviour is an often overlooked but important component of effective vector control. The collective results of this study indicate a potential role of the lure-and-kill approach to combat *L. infantum* transmission in Brazil. The protective effects were not dissimilar to those of the insecticide-impregnated collars, although the confidence intervals around all effect estimates were broad. Notwithstanding, it is reasonable to consider that robustly designed deployment of the lure-and-kill strategy could result in public and veterinary health benefits similar to those globally reported for the Scalibor collars. In order to maximise the synthetic pheromone efficacy, complimentary studies are underway to inform best practice for community-level deployment.

## Supporting information

## Acknowledgements

We especially thank the veterinary and entomological technicians for their unflagging efforts to complete the fieldwork, and A. Picado for helpful discussions. We are indebted to B. Krishnakumari for synthesis of the pheromone, to Russell IPM for preparing the slow release dispensers, to Ann Underhill for laboratory validation of the lure activity, and to S. Gokool for field assistance. V. Yardley supplied the initial *Leishmania* cultures, S. Mason enabled electronic cross-talk between laboratory and desktop hardware, and V. Camargo-Neves helped obtain historical canine data records. We also thank four external experts invited by the Wellcome Trust to review the trial design and statistical procedures. Finally, we acknowledge the public health workers and study communities for their compliance and participation.

## Authors’ Contributions

Conceptualization: Orin Courtenay, James G.C. Hamilton

Formal analysis: Orin Courtenay, Erin Dilger

Funding acquisition: James G.C. Hamilton, Orin Courtenay

Investigation: Erin Dilger, Leo A. Calvo-Bado, Lidija Kravar-Gard, Vicky Cooper, Melissa J. Bell, Raquel Goncalves, Muhammad M. Makhdoomi, Mikel A. González, Daniel P. Bray

Methodology: Erin Dilger, James G.C. Hamilton, Orin Courtenay

Project administration: Erin Dilger, Daniel P. Bray, Melissa J. Bell, Lidija Kravar-Gard, MJB, RPB, Caris M. Nunes, James G.C. Hamilton, Orin Courtenay

Resources: Reginaldo P. Brazil, Orin Courtenay, Caris M. Nunes

Supervision: Reginaldo P. Brazil, Caris M. Nunes, James G.C. Hamilton, Orin Courtenay

Writing-original draft: Orin Courtenay, James G.C. Hamilton, Erin Dilger

Writing-review & editing: James G.C. Hamilton, Orin Courtenay, Erin Dilger, Leo A. Calvo-Bado, Lidija Kravar-Gard, Vicky Cooper, Mikel A. González, Caris M. Nunes, Daniel P. Bray

## Funding

The work was funded by a Wellcome Trust Strategic Translation Award (WT091689MF) https://wellcome.ac.uk The funding body played no role in the design of the study, the collection, analysis, or interpretation of the data, or in writing the manuscript or decision to submit the paper for publication.

## Data sharing

The data supporting the conclusions of this article are included within the article and raw data will be available from the corresponding author upon reasonable request.

## Role of the funding source

The sponsors of the study and the companies providing the pheromone, insecticide and collars played no role in the design, data collection, data analysis, data interpretation, writing of the report, or decision to submit the paper for publication. The authors declare no commercial interests. The corresponding author confirm that they had full access to all the data in the study and had final responsibility for the decision to submit for publication.

## Consent for publication

Not applicable

## Availability of data and material

Data are available from the corresponding authors on reasonable request to be used solely within the context of this study, following the ethical agreements, and with permission from the relevant authorities and co-authors.

## Declaration of interests

The authors declare that they have no competing interests.

## Supplementary Information

S1. Intervention dates and intervals for the three trial arms.

S2. Numbers of dogs recruited to trial arms per period of the study.

S3. Laboratory methods

S4. Number of conversions to seropositive and parasite positive per cluster across strata and intervention arms amongst recruited dogs. Crude annual incidence shown for both measures.

S5. Summary of sand fly trapping effort and capture success in the trial arms.

## References

1. WHO. Global vector control response 2017–2030. Geneva: 2017 Contract No.: CC BY-NC-SA 3.0 IGO.

2. Bhatt S, Weiss DJ, Cameron E, Bisanzio D, Mappin B, Dalrymple U, et al. The effect of malaria control on Plasmodium falciparum in Africa between 2000 and Nature. 2015;526(7572):207-+. doi: 10.1038/nature15535. PubMed PMID: WOS:000362399000037.

3. Thomsen EK, Koimbu G, Pulford J, Jamea-Maiasa S, Ura Y, Keven JB, et al. Mosquito Behavior Change After Distribution of Bednets Results in Decreased Protection Against Malaria Exposure. Journal of Infectious Diseases. 2017;215(5):790–7. doi: 10.1093/infdis/jiw615. PubMed PMID: sWOS:000398636500020.

4. Shani A. Chemical communication agents (pheromones) in integrated pest management. Drug Development Research. 2000;50(3-4):400–5. doi: 10.1002/1098-2299(200007/08)50:3/4<400::aid-ddr22>3.0.co;2-v. PubMed PMID: WOS:000089628000022.

5. Cook SM, Khan ZR, Pickett JA. The use of push-pull strategies in integrated pest management. Annual Review of Entomology. 2007;52:375–400. doi: 10.1146/annurev.ento.52.110405.091407. PubMed PMID: WOS:000243653800019.

6. Mansour R, Belzunces LP, Suma P, Zappala L, Mazzeo G, Grissa-Lebdi K, et al. Vine and citrus mealybug pest control based on synthetic chemicals. A review. Agronomy for Sustainable Development. 2018;38(4). doi: 10.1007/s13593-018-0513-7. PubMed PMID: WOS:000438242300001.

7. Logan JG, Birkett MA. Semiochemicals for biting fly control: their identification and exploitation. Pest Management Science. 2007;63(7):647–57. doi: 10.1002/ps.1408. PubMed PMID: WOS:000247958400006.

8. Alvar J, Velez ID, Bern C, Herrero M, Desjeux P, Cano J, et al. Leishmaniasis Worldwide and Global Estimates of Its Incidence. Plos One. 2012;7(5). doi: 10.1371/journal.pone.0035671. PubMed PMID: WOS:000305338500009.

9. Quinnell RJ, Courtenay O. Transmission, reservoir hosts and control of zoonotic visceral leishmaniasis. Parasitology. 2009;136(14):1915–34. doi: 10.1017/s0031182009991156. PubMed PMID: WOS:000273515200006.

10. Alexander B, de Carvalho RL, McCallum H, Pereira MH. Role of the domestic chicken (Gallus gallus) in the epidemiology of urban visceral leishmaniasis in Brazil. Emerging Infectious Diseases. 2002;8(12):1480–5. doi: 10.3201/eid0812.010485. PubMed PMID: WOS:000179834300019.

11. Ministério da Saúde B. Manual de vigilância e controle da leishmaniose visceral In: Epidemiológica SdVeSDdV, editor. 1 ed. Brasília: Ministério da Saúde.; 2014. p. 120.

12. Camargo-Neves VLF, Glasser, C.M., Cruz, L.L., de Almeida, R.G. Manual de Vigilancia e Control da Leishmaniose Visceral Americana de Estado de Sao Paulo. Sao Paulo: 2006.

13. Bezerra JMT, de Araujo VEM, Barbosa DS, Martins-Melo FR, Werneck GL, Carneiro M. Burden of leishmaniasis in Brazil and federated units, 1990-2016: Findings from Global Burden of Disease Study 2016. Plos Neglected Tropical Diseases. 2018;12(9). doi: 10.1371/journal.pntd.0006697. PubMed PMID: WOS:000446054600011.

14. Shimozako HJ, Wu JH, Massad E. The Preventive Control of Zoonotic Visceral Leishmaniasis: Efficacy and Economic Evaluation. Computational and Mathematical Methods in Medicine. 2017. doi: 10.1155/2017/4797051. PubMed PMID: WOS:000402296600001.

15. PAHO/WHO. Leishmaniasis: Epidemiological Report in the Americas 2018; 6:[7 p.]. Available from: http://iris.paho.org/xmlui/handle/123456789/34856.

16. Seva AD, Mao L, Galvis-Ovallos F, Lima JMT, Valle D. Risk analysis and prediction of visceral leishmaniasis dispersion in Sao Paulo State, Brazil. Plos Neglected Tropical Diseases. 2017;11(2). doi: 10.1371/journal.pntd.0005353. PubMed PMID: WOS:000395741700028.

17. Harhay MO, Olliaro PL, Costa DL, Costa CHN. Urban parasitology: visceral leishmaniasis in Brazil. Trends in Parasitology. 2011;27(9):403–9. doi: 10.1016/j.pt.2011.04.001. PubMed PMID: WOS:000295207500007.

18. Krishnakumari B, Sarita Raj, K., Hamilton, J.G.C., editor Synthesis of 9-methylgermacrene from germacrone, an active analogue of (S)-9-methylgermacrene-B, sex pheromone of Phlebotomine sandfly, Lutzomyia longipalpis, from Lapinha Brazil. IUPAC International conference on Biodiversity and Natural Products: Chemistry and Medical Applications; 2004 26-31 January 2004; New Delhi (India).

19. Bray DP, Carter V, Alves GB, Brazil RP, Bandi KK, Hamilton JGC. Synthetic Sex Pheromone in a Long-Lasting Lure Attracts the Visceral Leishmaniasis Vector, Lutzomyia longipalpis, for up to 12 Weeks in Brazil. Plos Neglected Tropical Diseases. 2014;8(3). doi: 10.1371/journal.pntd.0002723. PubMed PMID: WOS:000337348800009.

20. Kelly DW, Dye C. Pheromones, kairomones and the aggregation dynamics of the sandfly Lutzomyia longipalpis. Animal Behaviour. 1997;53:721–31. doi: 10.1006/anbe.1996.0309. PubMed PMID: WOS:A1997WW45300005.

21. Morrison AC, Ferro C, Pardo R, Torres M, Wilson ML, Tesh RB. Nocturnal activity patterns of Lutzomyia longipalpis (Diptera, Psychodidae) at an endemic focus of visceral leishmaniasis in Colombia. Journal of Medical Entomology. 1995;32(5):605–17. doi: 10.1093/jmedent/32.5.605. PubMed PMID: WOS:A1995RT60800007.

22. Quinnell RJ, Dye C. An experimental study of the peridomestic distribution of Lyzomyia longipalpis (Diptera, Psychodidae). Bulletin of Entomological Research. 1994;84(3):379–82. PubMed PMID: WOS:A1994QA19700012.

23. Bell MJ, Sedda L, Gonzalez MA, de Souza CF, Dilger E, Brazil RP, et al. Attraction of Lutzomyia longipalpis to synthetic sex-aggregation pheromone: Effect of release rate and proximity of adjacent pheromone sources. Plos Neglected Tropical Diseases. 2018;12(12). doi: 10.1371/journal.pntd.0007007. PubMed PMID: WOS:000455103100037.

24. Bray DP, Bandi KK, Brazil RP, Oliveira AG, Hamilton JGC. Synthetic Sex Pheromone Attracts the Leishmaniasis Vector Lutzomyia longipalpis (Diptera: Psychodidae) to Traps in the Field. Journal of Medical Entomology. 2009;46(3):428–34. doi: 10.1603/033.046.0303. PubMed PMID: WOS:000265803800003.

25. Bray DP, Alves GB, Dorval ME, Brazil RP, Hamilton JGC. Synthetic sex pheromone attracts the leishmaniasis vector Lutzomyia longipalpis to experimental chicken sheds treated with insecticide. Parasites & Vectors. 2010;3. doi: 10.1186/1756-3305-3-16. PubMed PMID: WOS:000276434700001.

26. Camargo-Neves VLF. American Visceral Leishmaniasis in the state of São Paulo: current situation. Boletim Epidemiológico Paulista. 2007;4(48):12–4.

27. Cardim MFM, Rodas LAC, Dibo MR, Guirado MM, Oliveira AM, Chiaravalloti-Neto F. Introduction and expansion of human American visceral leishmaniasis in the state of Sao Paulo, Brazil, 1999-2011. Revista De Saude Publica. 2013;47(4):691–700. doi: 10.1590/s0034-8910.2013047004454. PubMed PMID: WOS:000328944200006.

28. Rangel O, HiramotoI, R.M., Henriques, L.d.F., Taniguchi, H.H., CiaravoloI, R.M.d.C., TolezanoI, J.E., França, A.C.C., Yamashiro, J., Oliveira, S.S.d. Epidemiological classification of cities according to the Program of Surveillance and Control of American Visceral Leishmaniasis in the State of São Paulo, updated in 2013. Boletim Epidemiológico Paulista. 2013;10(111):3–14.

29. CiaravoloI RMC, Oliveira, S.S., HiramotoI, R.M., Henriques, L.F., Taniguchi, H.H., Junior, A.V., Spinola, R., RangelI, O., Tolezano, J.E. Classificação Epidemiológica dos Municípios Segundo o Programa de Vigilância e Controle da Leishmaniose Visceral no Estado de São Paulo, dezembro de 2014. Boletim Epidemiologico Paulista. 2015;12(143):9–22.

30. Ministério de Saúde. Sistema de informacao de agravos de notificacao (SINAN). [Internet]. 2019 [cited 25/01/19]. Available from:http://portal.saude.sp.gov.br/cve-centro-de-vigilancia-epidemiologica-prof.-alexandre-vranjac/areas-de-vigilancia/doencas-de-transmissao-por-vetores-e-zoonoses/agravos/leishmaniose-visceral/dados-estatisticos.

31. Dye C, Davies CR, Lainson R. Communication among Phlebotomine sandflies - a field study of domesticated Lutzomyia longipalpis populations in Amazonian Brazil Animal Behaviour. 1991;42:183–92. doi: 10.1016/s0003-3472(05)80549-4. PubMed PMID: WOS:A1991GE61300002.

32. Galvis-Ovallos F, Casanova C, Bergamaschi DP, Galati EAB. A field study of the survival and dispersal pattern of Lutzomyia longipalpis in an endemic area of visceral leishmaniasis in Brazil. Plos Neglected Tropical Diseases. 2018;12(4). doi: 10.1371/journal.pntd.0006333. PubMed PMID: WOS:000433487700016.

33. Sidhu S. Evaluation of social and economic factors affecting implementation of novel vector control in Brazil: University of Warwick; 2010.

34. Paulin S, Frenais R, Thomas E, Baldwin PM. Laboratory assessment of the anti-feeding effect for up to 12 months of a slow release deltamethrin collar (Scalibor (R)) against the sand fly Phlebotomus perniciosus in dogs. Parasites & Vectors. 2018;11. doi: 10.1186/s13071-018-3094-z. PubMed PMID: WOS:000445964400002.

35. David JR, Stamm LM, Bezerra HS, Souza RN, Killick-Kendrick R, Lima JWO. Deltamethrin-impregnated dog collars have a potent anti-feeding and insecticidal effect on Lutzomyia longipalpis and Lutzomyia migonei. Memorias Do Instituto Oswaldo Cruz. 2001;96(6):839–47. PubMed PMID: WOS:000170361700018.

36. Killick-Kendrick R, Killick-Kendrick, M., Focheux, C., Dereure, J., Puech, M-P., Cadiergues, M.C. Protection of dogs from bites of phelbotomine sandflies by deltamethrin collars for control of canine leishmaniasis. Medical and Veterinary Entomology. 1997;(11):105–11.

37. Borja LS, de Sousa OMF, Solca MD, Bastos LA, Bordoni M, Magalhaes JT, et al. Parasite load in the blood and skin of dogs naturally infected by Leishmania infanturn is correlated with their capacity to infect sand fly vectors. Veterinary Parasitology. 2016;229:110–7. doi: 10.1016/j.vetpar.2016.10.004. PubMed PMID: WOS:000388549200019.

38. Da Silva VG. Aspectos entomológicos e infecção natural dos flebotomíneos por Leishmania (Leishmania) infantum chagasi em municípios do estado de São Paulo com autoctonia de transmissão de leishmaniose visceral humana e/ou canina [Mestre em Ciências]2016.

39. Hayes RJ, Moulton, L.H. Cluster randomized trials. Interdisciplinary Statistic Series. Florida: Chapman & Hall/CRC; 2009.

40. Hayes RJ, Bennett S. Simple sample size calculation for cluster-randomized trials. International Journal of Epidemiology. 1999;28(2):319–26. doi: 10.1093/ije/28.2.319. PubMed PMID: WOS:000079942800022.

41. Bottomley C, Kirby MJ, Lindsay SW, Alexander N. Can the buck always be passed to the highest level of clustering? Bmc Medical Research Methodology. 2016;16. doi: 10.1186/s12874-016-0127-1. PubMed PMID: WOS:000371566400001.

42. Naylor JC, Smith AFM. Applications of a method for the efficient computation of posterior distributions. Journal of the Royal Statistical Society Series C-Applied Statistics. 1982;31(3):214–25. PubMed PMID: WOS:A1982QC87400003.

43. Camargo-Neves V. Aspectos epidemiológicos e avaliação das medidas de controle da Leishmaniose Visceral Americana no Estado de São Paulo, Brasil.: Universidade de São Paulo - USP; 2004.

44. Dilger E. The effects of host-vector relationships and density dependence on the epidemiology of visceral leishmaniasis.: University of Warwick; 2013.

45. Brown MB, Forsythe AB. Robust test for equality of variances. Journal of the American Statistical Association. 1974;69(346):364–7. doi: 10.2307/2285659. PubMed PMID: WOS:A1974T799200012.

46. Courtenay O, Macdonald DW, Lainson R, Shaw JJ, Dye C. Epidemiology of canine leishmaniasis-a comparative serological study of dogs and foxes in Amazon Brazil. Parasitology. 1994;109:273–9. PubMed PMID: WOS:A1994PK02000002.

47. Antoniou M, Messaritakis I, Christodoulou V, Ascoksilaki I, Kanavakis N, Sutton AJ, et al. Increasing Incidence of Zoonotic Visceral Leishmaniasis on Crete, Greece. Emerging Infectious Diseases. 2009;15(6):932–4. doi: 10.3201/eid1506.071666. PubMed PMID: WOS:000266539100014.

48. Lira RA, Cavalcanti MP, Nakazawa M, Ferreira AGP, Silva ED, Abath FGC, et al. Canine visceral leishmaniosis: A comparative analysis of the EIE-leishmaniosevisceral-canina-Bio-Manguinhos and the IFI-leishmaniose-visceral-canina-Bio-Manguinhos kits. Veterinary Parasitology. 2006;137(1-2):11–6. doi: 10.1016/j.vetpar.2005.12.020. PubMed PMID: WOS:000236657200002.

49. Kazimoto TA, Amora SSA, Figueiredo FB, Magalhaes JME, Freitas YBN, Sousa MLR, et al. Impact of 4% Deltamethrin-Impregnated Dog Collars on the Prevalence and Incidence of Canine Visceral Leishmaniasis. Vector-Borne and Zoonotic Diseases. 2018;18(7):356–63. doi: 10.1089/vbz.2017.2166. PubMed PMID: WOS:000430622200001.

50. Reithinger R, Coleman PG, Alexander B, Vieira EP, Assis G, Davies CR. Are insecticide-impregnated dog collars a feasible alternative to dog culling as a strategy for controlling canine visceral leishmaniasis in Brazil? International Journal for Parasitology. 2004;34(1):55–62. doi: 10.1016/j.ipara.2003.09.006. PubMed PMID: WOS:000188377300007.

51. Lopes EG, Seva AP, Ferreira F, Nunes CM, Keid LB, Hiramoto RM, et al. Vaccine effectiveness and use of collar impregnated with insecticide for reducing incidence of Leishmania infection in dogs in an endemic region for visceral leishmaniasis, in Brazil. Epidemiology and Infection. 2018;146(3):401–6. doi: 10.1017/s0950268817003053. PubMed PMID: WOS:000424741000019.

52. Camargo-Neves V, Rodas LAC, Pauliquévis C Jr. Avaliação da efetividade da utilização de coleiras impregnadas com deltametrina a 4% para o controle da leishmaniose visceral americana no Estado de Sao Paulo: resultados preliminares. São Paulo: Boletim Epidemiológico Paulista. 2004;1(12):1–11.

53. Oliveira-Lima JW, Nonato de Souza, R., Teixeira, M. J., Pompeu, M., Killick-Kendrick, R., & David, J. R., editor Preliminary results of a field trial to evaluate deltamethrin-impregnated collars for the control of canine leishmaniasis in northeast Brazil. 2002; Seville, Spain: Intervet International bv.

54. Maroli M, Mizzoni V, Siragusa C, D’Orazi A, Gradoni L. Evidence for an impact on the incidence of canine leishmaniasis by the mass use of deltamethrin-impregnated dog collars in southern Italy. Medical and Veterinary Entomology. 2001;15(4):358–63. doi: 10.1046/j.0269-283x.2001.00321.x. PubMed PMID: WOS:000172903600002.

55. Foglia Manzillo V, Oliva G, Pagano A, Manna L, Maroli M, Gradoni L. Deltamethrin-impregnated collars for the control of canine leishmaniasis: evaluation of the protective effect and influence on the clinical outcome of Leishmania infection in kennelled stray dogs. Vet Parasitol. 2006;142(1-2):142–5. doi: 10.1016/j.vetpar.2006.06.029. PubMed PMID: 16884851.

56. Manzillo VF, Oliva G, Pagano A, Manna L, Maroli M, Gradoni L. Deltamethrin-impregnated collars for the control of canine leishmaniasis: Evaluation of the-protective effect and influence on the clinical outcome of Leishmania infection in kennelled stray dogs. Veterinary Parasitology. 2006;142(1-2):142–5. doi: 10.1016/j.vetpar.2006.06.029. PubMed PMID: WOS:000242232300017.

57. Ferroglio E, Poggi M, Trisciuoglio A. Evaluation of 65% permethrin spot-on and deltamethrin-impregnated collars for canine Leishmania infantum infection prevention. Zoonoses Public Health. 2008;55(3):145–8. doi: 10.1111/j.1863-2378.2007.01092.x. PubMed PMID: 18331517.

58. Aoun K, Chouihi, E., Boufaden, I., Mahmoud, R., Bouratbine, A., Bedoui, K. Efficacy of Deltamethrin-impreganated collars Scalibor in the prevention of canine leishmaniasis in the area of Tunis. Arch Inst Pasteur Tunis. 2008;85(1-4):63–8.

59. Gavgani ASM, Hodjati MH, Mohite H, Davies CR. Effect of insecticide-impregnated dog collars on incidence of zoonotic visceral leishmaniasis in Iranian children: a matched-cluster randomised trial. Lancet. 2002;360(9330):374–9. doi: 10.1016/s0140-6736(02)09609-5. PubMed PMID: WOS:000177255600011.

60. Courtenay O, Bazmani, A., Parvizi, P., Ready, P.D., Cameron, M.M. Insecticide–impregnated dog collars reduce infantile clinical visceral leishmaniasis under operational conditions in NW Iran: a community–wide cluster randomised trial. PLoS Neglected Tropical Diseases. 2019.

61. Silva RA, de Andrade AJ, Quint BB, Raffoul GES, Werneck GL, Rangel EF, et al. Effectiveness of dog collars impregnated with 4% deltamethrin in controlling visceral leishmaniasis in Lutzomyia longipalpis (Diptera: Psychodidade: Phlebotominae) populations. Memorias Do Instituto Oswaldo Cruz. 2018;113(5). doi: 10.1590/0074-02760170377. PubMed PMID: WOS:000428693000002.

62. Quinnell RJ, Dye C. Correlates of the peridomestic abundance of Lutzomyia longipalpis (Diptera, Psychodidae) in Amazon Brazil. Medical and Veterinary Entomology. 1994;8(3):219–24. doi: 10.1111/j.1365-2915.1994.tb00502.x. PubMed PMID: WOS:A1994NV98200003.

63. Kelly DW, Mustafa Z, Dye C. Differential application of lambda-cyhalothrin to control the sandfly Lutzomyia longipalpis. Medical and Veterinary Entomology. 1997;11(1):13–24. doi: 10.1111/j.1365-2915.1997.tb00285.x. PubMed PMID: WOS:A1997WK47000004.

64. Kelly DW, Mustafa Z, Dye C. Density-dependent feeding success in a field population of the sandfly, Lutzomyia longipalpis. Journal of Animal Ecology. 1996;65(4):517–27. doi: 10.2307/5786. PubMed PMID: WOS:A1996UZ02700011.

65. Jones TM, Quinnell RJ. Testing predictions for the evolution of lekking in the sandfly, Lutzomyia longipalpis. Animal Behaviour. 2002;63:605–12. doi: 10.1006/anbe.2001.1946. PubMed PMID: WOS:000175391400022.

66. Bray DP, Hamilton JGC. Host odor synergizes attraction of virgin female Lutzomyia longipalpis (Diptera: Psychodidae). Journal of Medical Entomology. 2007;44(5):779–87. doi: 10.1603/0022-2585(2007)44[779:hosaov]2.0.co;2. PubMed PMID: WOS:000249179000008.

67. Courtenay O, Quinnell RJ, Garcez LM, Shaw JJ, Dye C. Infectiousness in a cohort of Brazilian dogs: Why culling fails to control visceral leishmaniasis in areas of high transmission. Journal of Infectious Diseases. 2002;186(9):1314–20. doi: 10.1086/344312. PubMed PMID: WOS:000178577500014.

68. Courtenay O, Carson C, Calvo-Bado L, Garcez LM, Quinnell RJ. Heterogeneities in Leishmania infantum Infection: Using Skin Parasite Burdens to Identify Highly Infectious Dogs. Plos Neglected Tropical Diseases. 2014;8(1). doi: 10.1371/journal.pntd.0002583. PubMed PMID: WOS:000337977300007.

69. Vercosa BLA, Lemos CM, Mendonca IL, Silva S, de Carvalho SM, Goto H, et al. Transmission potential, skin inflammatory response, and parasitism of symptomatic and asymptomatic dogs with visceral leishmaniasis. Bmc Veterinary Research. 2008;4. doi: 10.1186/1746-6148-4-45. PubMed PMID: WOS:000262310000001.

70. de Amorim IFG, da Silva SM, Figueiredo MM, Moura EP, de Castro RS, Lima TKD, et al. Toll Receptors Type-2 and CR3 Expression of Canine Monocytes and Its Correlation with Immunohistochemistry and Xenodiagnosis in Visceral Leishmaniasis. Plos One. 2011;6(11). doi: 10.1371/journal.pone.0027679. PubMed PMID: WOS:000298168100015.

71. Laurenti MD, Rossi CN, da Matta VLR, Tomokane TY, Corbett CEP, Secundino NFC, et al. Asymptomatic dogs are highly competent to transmit Leishmania (Leishmania) infantum chagasi to the natural vector. Veterinary Parasitology. 2013;196(3-4):296–300. doi: 10.1016/j.vetpar.2013.03.017. PubMed PMID: WOS:000323864500008.

72. Giraud E, Martin, O., Yakob, L., Rogers, M. Quantifying Leishmania Metacyclic Promastigotes from Individual Sandfly Bites Reveals the Efficiency of Vector Transmission. Communications Biology. 2019;In Press.

73. Doehl JSP, Bright Z, Dey S, Davies H, Magson J, Brown N, et al. Skin parasite landscape determines host infectiousness in visceral leishmaniasis. Nature Communications. 2017;8. doi: 10.1038/s41467-017-00103-8. PubMed PMID: WOS:000404778800001.

74. Costa DN, Codeco CT, Silva MA, Werneck GL. Culling dogs in scenarios of imperfect control: realistic impact on the prevalence of canine visceral leishmaniasis. PLoS Negl Trop Dis. 2013;7(8):e2355. doi: 10.1371/journal.pntd.0002355. PubMed PMID: 23951375; PubMed Central PMCID: PMC3738479.

75. Travi BL, Cordeiro-da-Silva A, Dantas-Torres F, Miro G. Canine visceral leishmaniasis: Diagnosis and management of the reservoir living among us. Plos Neglected Tropical Diseases. 2018;12(1). doi: 10.1371/journal.pntd.0006082. PubMed PMID: WOS:000424022700011.

76. Dantas-Torres F, Otranto D. Best Practices for Preventing Vector-Borne Diseases in Dogs and Humans. Trends in Parasitology. 2016;32(1):43–55. doi: 10.1016/j.pt.2015.09.004. PubMed PMID: WOS:000368206900009.

